# A statistical framework for the robust detection of hidden variation in single cell transcriptomes

**DOI:** 10.1101/151217

**Authors:** Donghyung Lee, Anthony Cheng, Mohan Bolisetty, Duygu Ucar

## Abstract

Single cell RNA-sequencing (scRNA-seq) precisely characterize gene expression levels and dissect variation in expression associated with the state (technical or biological) and the type of the cell, which is averaged out in bulk measurements. Multiple and correlated sources contribute to gene expression variation in single cells, which makes their estimation difficult with the existing methods developed for bulk measurements (e.g., surrogate variable analysis (SVA)) that estimate orthogonal transformations of these sources. We developed iteratively adjusted surrogate variable analysis (IA-SVA) that can estimate hidden and correlated sources of variation by identifying a set of genes affected with each hidden factor in an iterative manner. Analysis of scRNA-seq data from human cells showed that IA-SVA could accurately capture hidden variation arising from technical (e.g., stacked doublet cells) or biological sources (e.g., cell type or cell-cycle stage). Furthermore, IA-SVA delivers a set of genes associated with the detected hidden source to be used in downstream data analyses. As a proof of concept, IA-SVA recapitulated known marker genes for islet cell subsets (e.g., alpha, beta), which improved the grouping of subsets into distinct clusters. Taken together, IA-SVA is an effective and novel method to dissect multiple and correlated sources of variation in scRNA-seq data.

## Introduction

Single-cell RNA-Sequencing (scRNA-seq) enable precise characterization of gene expression levels, which harbour variation in expression associated with both technical (e.g., biases in capturing transcripts from single cells, PCR amplifications or cell contamination) and biological sources (e.g., differences in cell cycle stage or cell types). If these sources are not accurately identified and properly accounted for, they might confound the downstream analyses and hence the biological conclusions^1-3^. In bulk measurements, hidden sources of variation are typically unwanted (e.g., batch effects) and are computationally eliminated from the data. However, in single cell RNA-seq data, variation/heterogeneity stemming from hidden biological sources can be the primary interest of the study; which necessitate their accurate detection (i.e., existence of a hidden factor) and estimation (i.e., contribution of this factor to the gene expression levels) for downstream data analyses and interpretation. For example, a recent such study uncovered a CD1C+ dendritic cell (DC) subset by profiling human blood samples^4^ and improved the immune monitoring of human DCs in health and disease. One challenge in detecting hidden sources of variation in scRNA-seq data lies in the existence of multiple and highly correlated hidden sources, including geometric library size (i.e., the library size of log-transformed read counts), number of expressed genes in a cell, experimental batch effects, cell cycle stage and cell type^5-8^. The correlated nature of hidden sources limits the efficacy of existing algorithms to accurately detect the source and estimate its contribution to the variation in the data.

‘Surrogate variable analysis’ (SVA)^9-11^ is a family of algorithms that are developed to detect and remove hidden and “unwanted” variation (e.g., batch effect) in gene expression data by accurately parsing the data into signal and noise. A number of SVA-based methods have been developed and used for the analyses of microarray, bulk, and single-cell RNA-seq data including SSVA^11^ (supervised surrogate variable analysis), USVA^10^ (unsupervised SVA), ISVA^12^ (Independent SVA), RUV (removing unwanted variation)^13,14^, and most recently scLVM^6^ (single-cell latent variable model). These methods primarily aim to remove ‘unwanted’ variation (e.g., batch or cell-cycle effect) in data while preserving the biological signal of interest typically to improve downstream differential expression analyses between cases and controls. For this purpose, they utilize PCA (principal component analysis), SVD (singular vector decomposition) or ICA (independent component analysis) to infer orthogonal transformations of hidden factors that can be used as covariates in downstream analysis. However, this paradigm by definition results in orthogonality between multiple estimated factors and limits the efficacy of existing SVA-based methods for single-cell data analyses, in which some of the sources of variation are ‘wanted’ and are highly correlated with each other.

To fill this gap, we developed a robust and iterative SVA-based statistical framework: **I**teratively **A**djusted **S**urrogate **V**ariable **A**nalysis (IA-SVA) (Figure 1A and Methods for details), which provides three major advantages. First, it accurately estimates multiple hidden sources of variation even if the sources are correlated with each other and with known sources, which is a limitation of existing SVA-based methods. Second, it enables assessing the significance of each detected factor for explaining the unmodeled variation in the data. Third, it delivers a set of genes that are significantly associated with the detected hidden source. Application of IA-SVA for scRNA-seq data analyses is diverse including the detection of “unwanted” variation due to cell contamination or “wanted” variation associated with rare cell types (Figure 1B). In simulation studies we showed that IA-SVA i) provides high statistical power in detecting hidden factors; ii) controls Type I error rate at the nominal level (α = 0.05); iii) delivers high accuracy in estimating hidden factors. We evaluated the efficacy of IA-SVA on scRNA-seq data from human pancreatic islets and brain cells and showed that IA-SVA is effective in capturing heterogeneity associated with both technical (e.g., doublet cells) and biological sources (e.g., differences in cell types or cell-cycle stages). Furthermore, we showed that IA-SVA based gene selection can be further utilized in downstream analyses such as in data visualization using t-distributed stochastic neighbor embedding (tSNE) ^15^ and performs favourably compared to existing methods developed for gene selection and visualization (e.g., Spectral tSNE^16^).

**Figure 1.**
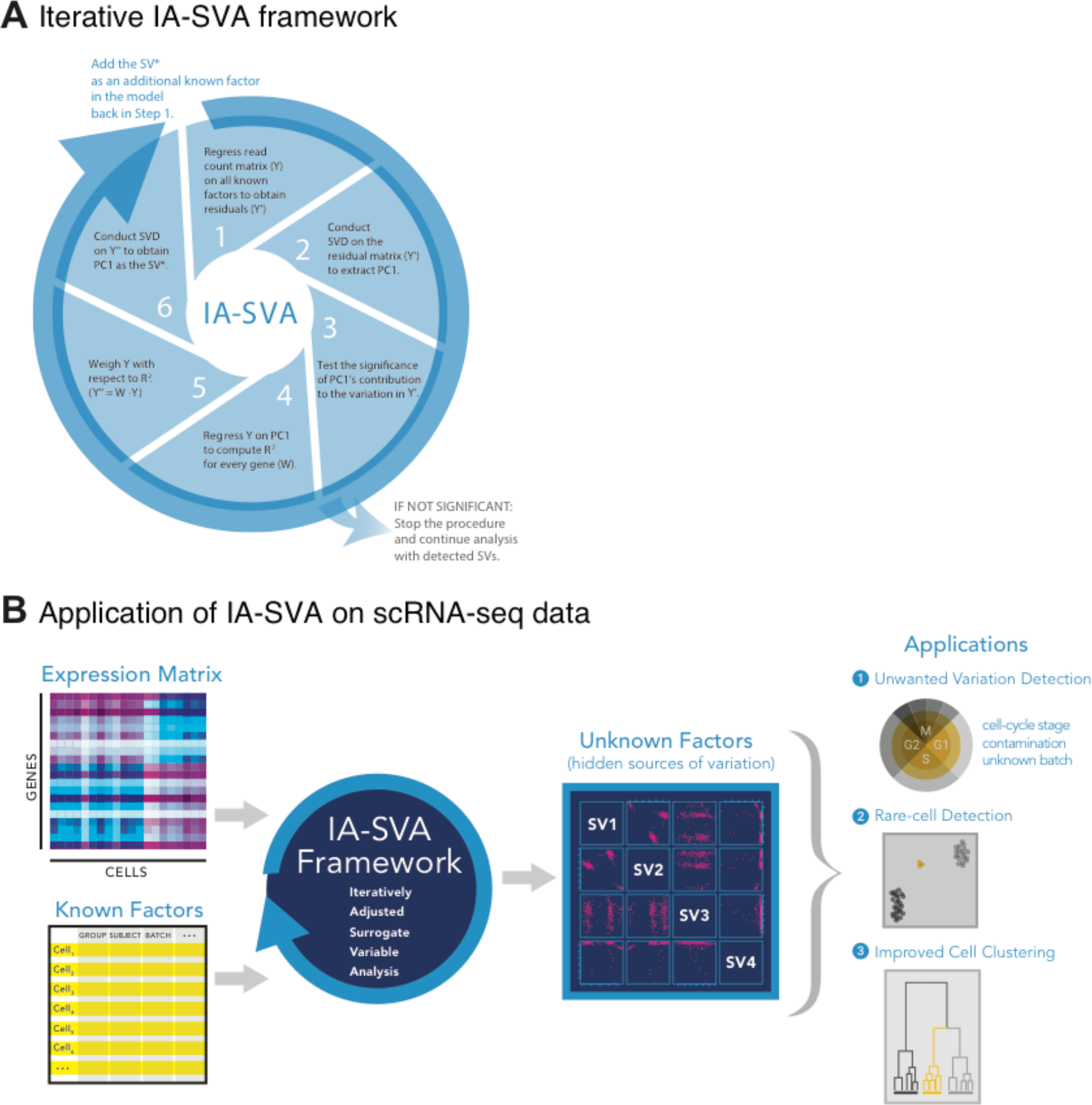
IA-SVA is a robust statistical framework to detect and estimate multiple and correlated hidden sources of variation. **(A)** Six-step IA-SVA procedure. IA-SVA computes the first principal component (PC1) from read counts adjusted for all known factors and tests its significance [Steps 1-3]. If significant, IA-SVA uses this PC1 to infer a set of genes associated with the hidden factor [Steps 4-5] and obtain a surrogate variable (SV) to represent the hidden factor using these genes [Step 6]. **(B)** IA-SVA uses single-cell gene expression data matrix and known factors to detect hidden sources of variation (e.g., cell contamination, cell-cycle status, and cell type). If these factors match to a biological variable of interest (e.g., cell type assignment), genes highly correlated with the factor can be detected and used in downstream analyses (e.g., data visualization).

## Results

### Benchmarking IA-SVA on simulated data

To assess and compare the detection power, Type I error rate, and the accuracy of hidden source estimates using IA-SVA and existing state-of-the-art methods (i.e., USVA and SSVA), we performed simulation studies (see Methods for details) under the null hypothesis (i.e., a group (case/control) variable affecting 10% of genes and no hidden factor) and under the alternative hypothesis (i.e., a group variable and three hidden factors affecting 30%, 20%, 10% of genes, respectively). Under the alternative hypothesis, we considered two correlation scenarios where the three hidden factors are moderately (|*r*|=~0.3-0.6) or weakly (|*r*|<0.3) correlated with the group variable (i.e., a known factor). Under each simulation scenario, we generated 1,000 scRNA-seq data sets (10,000 genes and 50 cells) and performed IA-SVA, USVA and SSVA (α =0.05, 50 permutations) on them to detect simulated hidden factors. Using these simulation results, we assessed the empirical Type I error rate of each method (i.e., the number of times each method detects a false positive factor under the null hypothesis at the nominal level of 0.05 divided by the number of simulations (n=1,000)). Similarly, we also quantified the empirical detection power rate of each method under different alternative hypothesis scenarios as the number of times each method detects a simulated factor under the alternative hypothesis (i.e., a factor actually exists and is detected as significant by the method) divided by the number of simulations. We used the average of the absolute Pearson correlation coefficients between the simulated and estimated hidden factors to quantify the accuracy of estimates.

Simulation studies showed that IA-SVA performs as well or better than USVA and SSVA in terms of detection power and accuracy of the estimate while controlling the Type I error rate (0.04 for IA-SVA versus 0.09 for USVA and SSVA) (Table 1). In particular, IA-SVA was more effective when a hidden factor affected a small percentage of genes and when the factors were correlated (|*r*|=0.3-0.6) with the known factor (i.e., group variable). For example IA-SVA detected Factor3, which affected only 10% genes, 87% of the time, whereas USVA and SSVA detected this factor 78% of the simulations (first three columns in Table 1). More importantly, IA-SVA correctly inferred the correlations among multiple hidden factors while USVA and SSVA delivered biased estimates due to their orthogonality assumption (Supplementary Figure S1).

**Table 1.**
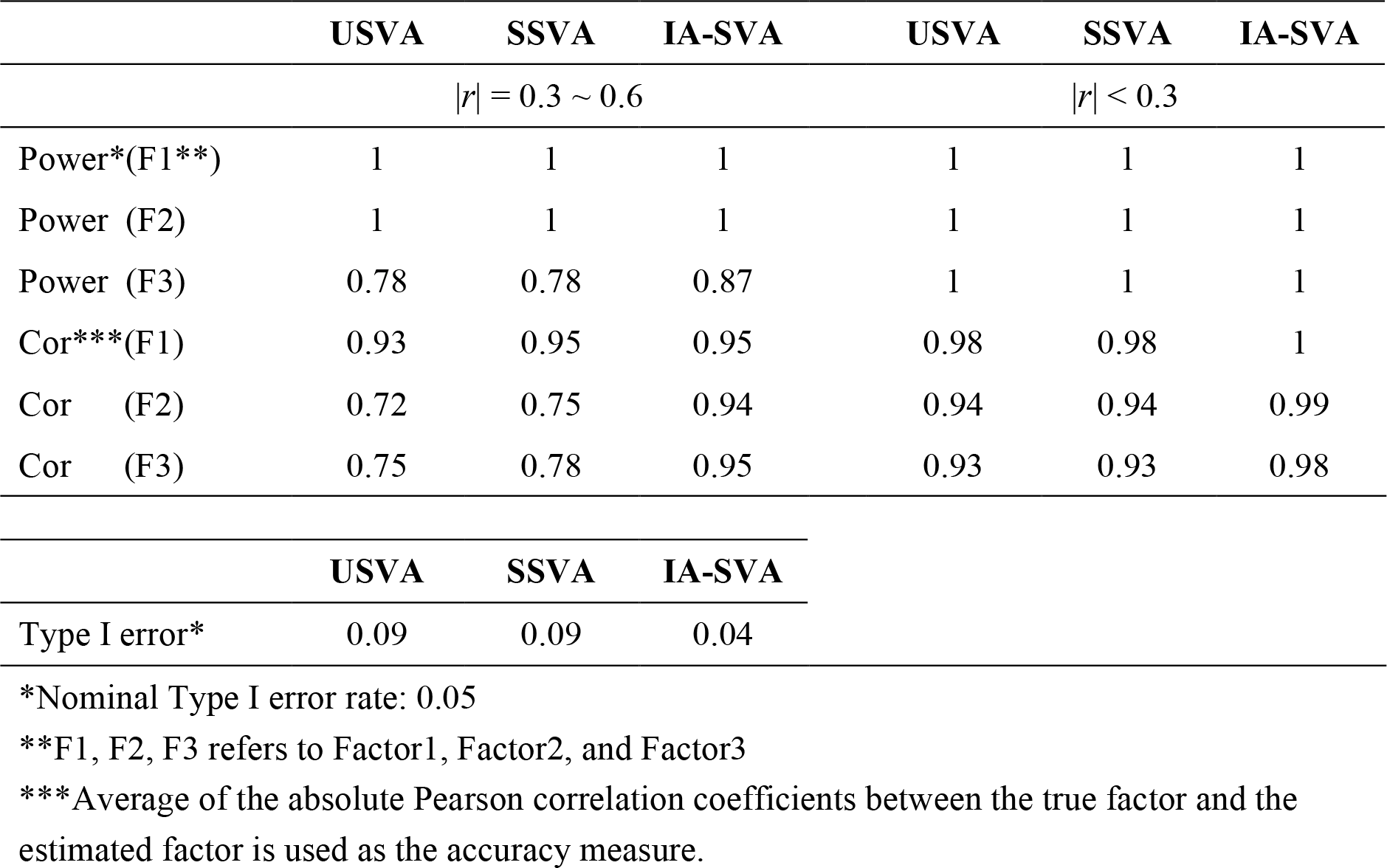
IA-SVA accurately captures unknown sources of variation while controlling Type I error rate at a nominal level. Empirical power, Type I error rate, and the accuracy of estimates for IA-SVA, SSVA, and USVA assessed using simulated single-cell gene expression data. Alternative scenarios are simulated in which hidden factors are moderately (|*r*|*=~*0.3-0.6, first three columns) or weakly (|*r*|*<*0.3, last three columns) correlated with the group variable. IA-SVA outperforms alternative methods especially while detecting variation stemming from a smaller fraction of genes (10%) and especially when factors are correlated.

### IA-SVA captures variation stemming from a small number of alpha cells

To test whether IA-SVA is effective in capturing variation within a homogenous cell population, we analysed scRNA-seq data generated from human alpha cells (n=101, marked with glucagon (*GCG*) expression) obtained from three diabetic patients^17^ using the Fluidigm C1 platform^18^, for which the original study did not report any separation of these alpha cells. Using geometric library size and patient ID as known factors, significant surrogate variables (SVs) were inferred using IA-SVA (α =0.05, 50 permutations) on the data (14,416 genes and 101 cells). For comparison, we applied PCA, USVA, and tSNE on this data. In USVA analysis, similarly geometric library size and patient ID were used as known factors and significant SVs were obtained (α =0.05, 50 permutations). In the PCA analysis, PC1 was discarded since it is highly correlated (*r* = 0.99) with the geometric library size.

Top two significant SVs inferred by IA-SVA clearly separated alpha cells into two groups (six outlier cells marked in red vs. the rest marked in grey at SV2 > 0.1) (Figure 2A). 27 genes significantly associated with second SV (SV2) (Benjamini-Hochberg q-value (FDR) < 0.05, coefficient of determination (*R^2^*) > 0.6), which included genes expressed in fibroblasts such as *COL4A1* and *COL4A2*. These genes were exclusively expressed in six outlier cells and clearly separated alpha cells into two clusters (Figure 2B). A larger set of SV2-associated genes (n = 108, FDR < 0.05, *R^2^* > 0.3) was used for pathway and GO enrichment analyses and uncovered that these genes are associated with extracellular matrix receptors (Supplementary Table S1). Hence, these outlier cells likely arise from cell contamination (e.g., fibroblasts contaminating islet cells) or cell doublets (e.g., two cells captured together) — a known problem in early Fluidigm C1 experiments^20,21^. Alternative methods (i.e., PCA, USVA, tSNE) failed to clearly detect these outlier cells (Figure 2C-E).

**Figure 2.**
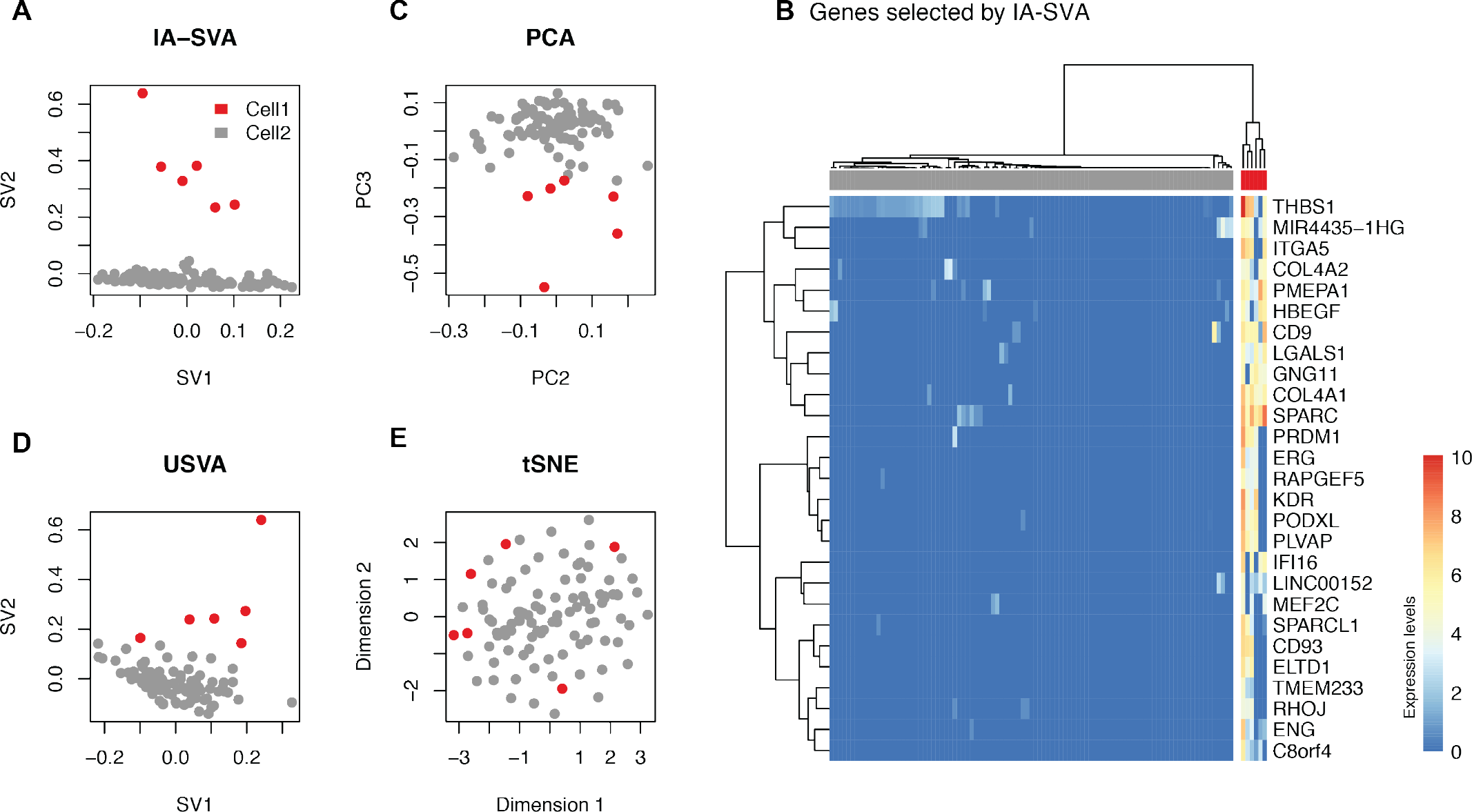
IA-SVA can detect heterogeneity originating from potentially contaminated alpha cells. (**A**) Outlier alpha cells captured using IA-SVA and same cells marked in respective (**C**) PCA, (**D**) USVA, and (**E**) tSNE analyses. Cells are clustered into two groups (red vs. gray dots) based on IA-SVA’s surrogate variable 2 (SV2 > 0.1). In PCA, PC1 was discarded since it explains the geometric library size. (**B**) Hierarchical clustering of alpha cells using 27 genes significantly associated with SV2 (FDR < 0.05 and *R^2^* > 0.6) (ward.D2 and cutree_cols =2). 6 cells clearly separated from the rest of the alpha cells based on the expression of these 27 genes.

We next studied whether this source of heterogeneity can be recapitulated in an independent and bigger human islet scRNA-seq dataset^18^, using gene expression profiles (17,168 genes) of 569 alpha cells from six diabetic patients. Using geometric library size and patient ID as known factors we identified top 2 significant SVs using IA-SVA and USVA. For comparison, we also conducted PCA and tSNE analyses on this data. In PCA, PC1 was discarded since it matched the geometric library size, which is adjusted for in IA-SVA and USVA analyses. IA-SVA’s SV2 separated alpha cells into two groups (Supplementary Figure S2A) and as in the previous case it was associated with fibrotic response genes including *SPARC, COL4A1, COL4A2* (n=81, FDR < 0.05 and *R^2^* > 0.3) (Supplementary Figure S2B, GO/pathway results in Supplementary Table S2). These results highlight IA-SVA’s ability to detect variation among alpha cells potentially due to cell contamination or cell doublets. PCA, USVA, and tSNE failed to clearly separate these compromised alpha cells (Supplementary Figure S2C-E) from the rest of the cells.

### IA-SVA accurately detects variation arising from cell-cycle stage differences

Differences in cell-cycle stages lead to variation in single cell gene expression data^3^. Supervised methods based on SVA have been developed to detect and correct for cell cycle stage differences, most notably the scLVM algorithm. scLVM implements a Bayesian latent variable model to infer hidden cell-cycle factors by using known cell cycle genes^6^. IA-SVA can provide an unsupervised alternative by accurately capturing cell-cycle related variation in single cell data. To show this, we analyzed scRNA-seq data (21,907 genes and 74 cells) obtained from human glioblastomas that has an established cell-cycle signature^22^. We conducted IA-SVA analyses by using geometric library size as a known factor and extracted top 2 significant SVs (α=0.05, 50 permutations). For comparison, we applied PCA, USVA and tSNE analyses on this data, where for USVA geometric library size is used as a known factor.

IA-SVA’s SV1 clearly separated 13 cells from the rest (cells marked in red in Figure 3A), which was associated with 119 genes (FDR < 0.05 and *R^2^* > 0.3). Hierarchical clustering (ward.D2, cutree_cols=2) using these genes confirmed the separation of cells into two groups (Figure 3B), whereas alternative methods failed to clearly separate these two groups of cells (Figure 3C-E). Pathway and GO enrichment analyses of these genes^23,24^ revealed significant enrichment for cell-cycle process related GO terms and pathways (Supplementary Table S3), suggesting that this hidden variation is stemming from cell-cycle stage differences. Indeed, cell-cycle-stage predictions of cells using the SCRAN R package^25^ showed that cells in different cell-cycle stages have different SV1 values (Figure 3F). We noted that SV1 is highly correlated (|*r*|= 0.44) with the geometric library size (typically the top contributor to the variation in single cell data), which might explain why alternative methods failed to clearly detect this variation in the data. These results demonstrate that IA-SVA can effectively detect variation stemming from cell-cycle differences in an unsupervised manner from single cell transcriptomes, even if this factor is highly correlated with known factors.

**Figure 3.**
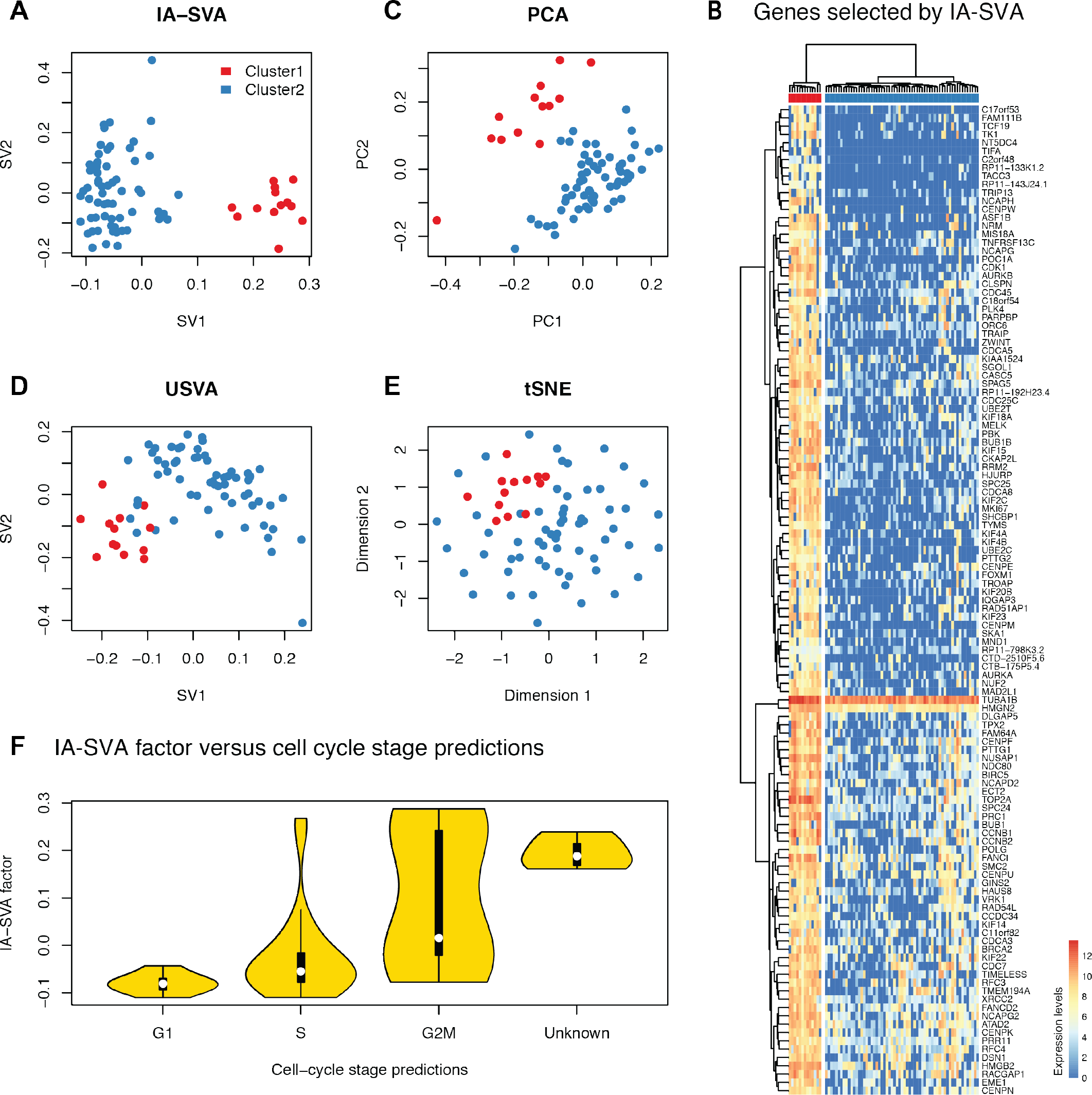
IA-SVA can detect heterogeneity stemming from differences in cell-cycle stage. (**A**) Visualization of glioblastoma cells based on IA-SVA-detected factors (SV1 and SV2). Same cells are marked in respective analyses with (**C**) PCA, (**D**) USVA, and (**E**) tSNE analyses. IA-SVA’s SV1 clearly separates cells into two groups (red vs. blue dots, SV1 > 0.1). Other methods failed to clearly detect this cell-cycle related heterogeneity. (**B**) Hierarchical clustering on 119 genes significantly associated (FDR < 0.05 and *R^2^* > 0.3) with IA-SVA’s SV2 confirms the separation of cells based on these genes (ward.D2 and cutree_cols = 2). (**F**) IA-SVA’s SV1 can segregate cells based on their cell-cycle-stage as predicted by SCRAN.

### IA-SVA based gene selection improves single cell data visualization

tSNE and other dimension reduction algorithms (e.g., Spectral tSNE implemented in Seurat^16^) are frequently used to visualize single cell data since they group together cells with similar gene expression patterns. However, variation introduced by technical or biological factors can confound the signal of interest and generate spurious clustering of data. IA-SVA can be particularly effective in handling this problem by estimating hidden factors of interest accurately while adjusting for all known factors of no interest. Moreover, IA-SVA identifies genes associated with each detected hidden factor, which could be biologically relevant such as marker genes for different cell types. The genes inferred by IA-SVA can significantly improve the performance of data visualization methods (e.g., tSNE^15^). To illustrate this, we studied single cell gene expression profiles (16,005 genes) of alpha (n=101, marked with glucagon (*GCG*) expression), beta (n=96, marked with insulin (*INS*) expression), and ductal (n=16, marked with *KRT19* expression) cells obtained from three diabetic patients^17^. First, we applied tSNE on all genes (n=16,005) and color-coded genes based on the reported cell type assignments^17^, which failed to separate cells from different origins (Figure 4A). Next, we applied IA-SVA on this data using patient ID, batch ID and the number of expressed genes as known factors and obtained significant SVs. SV1 and SV2 separated cells into distinct clusters (Supplementary Figure S3), suggesting that these SVs might be associated with cell type differences. Indeed, genes associated with SV1 and SV2 (n=92, FDR < 0.05 and R^2^ > 0.5) included known marker genes used in the original study (*INS*, *GCG*, *KRT19*) and uncovered alternative marker genes associated with alpha, beta and ductal cells (Figure 4B). These genes were annotated with diabetes and insulin processing related GO terms and pathways (Supplementary Table S4). As expected, tSNE analyses based on these 92 genes improved data visualization and clearly grouped together cells with respect to their cell type assignments (Figure 4C). Such improved analyses can be instrumental in discovering cells that might be incorrectly labelled based on a single marker gene. For example, our analyses revealed a beta cell that is labelled as a ductal cell in the original study (one green cell clustered with blue cells in Figure 4C). For comparison, we applied recently developed visualization methods, CellView^26^ and Spectral tSNE^16^, on the same data with their recommended settings. CellView identified the 1000 most over-dispersed genes and conducted tSNE on these genes. Spectral tSNE detected 2,933 most over-dispersed genes and performed tSNE on significant principal components of these genes. On this small dataset, both methods managed to group cells of different types into distinct groups (Supplementary Figure S4), suggesting that existing methods for gene selection and visualization are effective when datasets are small in size and are not confounded with multiple factors.

**Figure 4.**
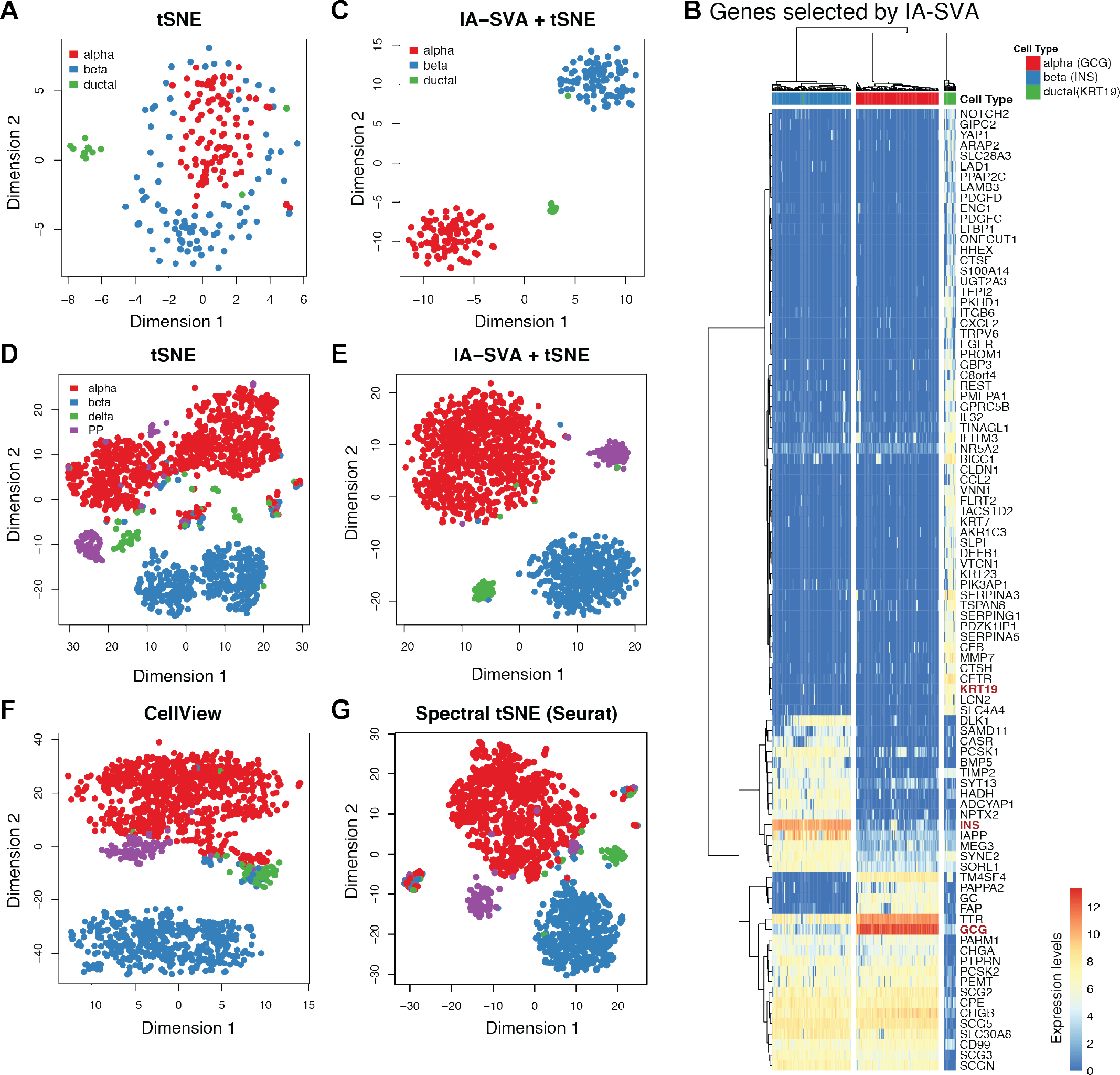
IA-SVA based gene selection enhances single cell data visualization. (**A**) tSNE analyses using all expressed genes in human islet data (tSNE). Cells are color-coded based on the original cell-type assignments. Note that cells are not effectively clustered with respect to their assigned cell types. (**B**) Hierarchical clustering using genes (n=92) selected by IA-SVA clearly separate cell types (ward.D2 and cutree_cols=3). Known marker genes (e.g., *INS*) are highlighted in red color. (**C**) tSNE analyses using the 92 IA-SVA genes (IA-SVA+tSNE). Note the improved grouping of cell types into discrete clusters. (**D**) tSNE analyses using top variable genes in a second and bigger islet scRNA-seq data. Note that cells are not effectively clustered with respect to their assigned cell types just using tSNE. (**E**) tSNE analyses repeated using genes (n=57) obtained via IA-SVA (IA-SVA+tSNE). Note the improved clustering of different cell types into discrete clusters. (**F**) tSNE analyses using 1000 most over-dispersed genes (CellView). (**G**) tSNE analyses on significant PCs obtained from highly over-dispersed genes (Spectral tSNE).

To test the efficacy of these methods on a bigger and more complex dataset, we conducted similar analyses on scRNA-seq data (19,226 genes) of 1,600 islet cells including alpha (n=946), beta (n=503), delta (n=58), and PP (n=93) cells from 6 diabetic and 12 non-diabetic individuals, where the study includes multiple confounding factors (e.g., ethnicity, disease state)^18^. We noted that original cell type assignments significantly correlate with patient identifications (*C*=0.48, *C*=Pearson’s contingency coefficient) and with ethnicity (*C*=0.25), which would reduce the ability of existing methods to detect variation associated with cell types. In such complex datasets, failing to properly adjust for potential confounding factors prior to data analyses can lead to spurious grouping of cells, which might mislead the biological conclusions. Indeed, when these cells were visualized using tSNE using all genes (n=19,226) and were color-coded with respect to the original cell-type assignments ^18^, cell types did not separate from each other and spurious clusters were observed within each cell type (Figure 4D). As suspected, potential confounding factors (i.e., patient ID and ethnicity) explained this grouping of cells (Supplementary Figure S5), which might be misleading as researchers are looking for alpha and beta cell subtypes that can be related to Type 2 Diabetes pathogenesis^27^. To eliminate spurious clusters stemming from known factors, existing methods (e.g., Seurat) simply regress out all known factors prior to visualization. However, this might affect the signal of interest (i.e., cell type assignment), due to high correlation between known factors (i.e., patient ID) and the hidden factor (i.e., cell types).

We applied IA-SVA on this complex data, while accounting for known factors (i.e., the number of expressed genes and patient ID) and extracted top four significant SVs (Supplementary Figure S6A and B). We identified 57 genes associated with the most significant SV (SV1) (FDR < 0.05 and *R^2^* > 0.5), which included known marker genes (i.e., *INS* and *GCG*) (Supplementary Figure S7, Supplementary Table S5) and revealed novel marker genes for these cell types. tSNE analyses using these 57 IA-SVA detected genes clearly separated different cell types into discrete groups and reinforced the importance of properly adjusting for known factors prior to data analyses (Figure 4E). For comparison, we applied CellView and Spectral tSNE on this data with recommended settings; however they failed to accurately group cells into distinct cell types (Figure 4F and G). Similar analyses were conducted using PCA and USVA on the same data, where top surrogate factors obtained with both methods failed to separate different cell types into distinct groups (Supplementary Figure S6C and D). Combined together these analyses suggest that IA-SVA is particularly effective in the analyses of complex datasets, which include the measurements of many cells that are affected by diverse confounding factors.

## Discussion

Surrogate variable analyses based methods are effective in detecting and eliminating hidden and unwanted variation in bulk gene expression data (such as batch effects). By using dimensionality reduction algorithms (e.g., PCA or SVD), these methods infer linear transformations of hidden factors and utilize these factor estimates as additional covariates in downstream analyses to eliminate unwanted variation^14^. However, measurement of gene expression levels at single cell resolution pose novel challenges in the detection and adjustment of hidden sources of data variation. First, single cell transcriptomes harbor hidden variation that can be biologically interesting (hence ‘wanted’) and can be the major goal of the study, for example detection of rare cells within a tissue^28^ or detection of a cell’s subtypes that can be linked to health or disease^27^. Second, since single cell data do not average out variation as in the case of bulk profiling, the data reflect variation arising from diverse biological and technical sources some of which could be highly correlated. Existing SVA-based methods do not readily apply to the unique needs of single cell data analyses. To fill this gap, we developed IA-SVA, where the objective is the accurate estimation of hidden factors even if these factors are correlated with each other or with the known factors. Unlike other SVA-based methods, IA-SVA focuses more on the accurate detection and estimation of hidden factors rather than their elimination since these factors can be biologically interesting, e.g., identification of a new cell type and its marker genes. Indeed, analyses on simulated scRNA-seq data showed that IA-SVA outperforms existing supervised (i.e., SSVA) and unsupervised (i.e., USVA) state-of-the-art methods in the estimation of hidden factors (not necessarily in their elimination). Furthermore, we noted that IA-SVA is particularly effective (i.e., high detection power and accuracy, and Type I error rate controlled under the nominal level of 0.05) in detecting correlated factors that affect a small fraction of genes. Therefore IA-SVA is an effective unsupervised alternative to existing SVA-based algorithms when the goal is to accurately estimate hidden factors (and their marker genes) rather than to eliminate these factors.

Through analyses of diverse human datasets from multiple studies, we established that IA-SVA can effectively detect hidden heterogeneity in scRNA-seq data arising from a small number of cells either due to technical (i.e., contamination or doublets) or biological (i.e., a rare cell type) sources. In two independently generated islet scRNA-seq datasets, we showed that IA-SVA detects heterogeneity stemming from compromised alpha cells (contaminated or stacked), which should be excluded from the downstream analyses (Figure 2 and Supplementary Figure S2). Therefore, IA-SVA provides an easy-to-apply statistical framework to uncover variation in scRNA-seq data even if it is stemming from only a handful of cells. This ability of IA-SVA can be effective in identifying rare cells within a population of cells, where genes associated with the detected factor can uncover relevant marker genes for the rare population of cells. In addition, IA-SVA can be effective in detecting heterogeneity associated with cell-cycle stages without prior knowledge, therefore providing an unsupervised solution to this common problem in single cell data analyses (Figure 3).

An important feature of IA-SVA is its ability to uncover genes associated with detected hidden factors. This feature can be used to detect marker genes associated with different cell types. As a proof-of-concept we demonstrated this in pancreatic islet cells, where we captured known marker genes (e.g., *INS, GCG*) in an unsupervised manner. Moreover, genes captured by IA-SVA can be used to improve the visualization of single cells into their respective clusters, as demonstrated with the analyses of islet cells from two separate studies (Figure 4). Spectral tSNE^16^ is a commonly used method for scRNA-seq data visualization especially in the existence of confounding factors. This method regresses out variation associated with known factors before data visualization. However, when a hidden factor is ‘wanted’ (e.g., cell types) and is highly correlated with known factors, removing the known factors will also diminish the ability to detect the wanted hidden factor and the genes associated with this factor (e.g., marker genes for different cell types). Indeed, our analyses using islet cells emphasized the importance of properly adjusting the data for known factors prior to further analyses, such as data visualization (e.g., tSNE) to prevent spurious clustering of cells due to the confounding factors (Figure 4E). IA-SVA is an alternative method that can effectively handle data with multiple confounding factors.

In summary, IA-SVA is an SVA-based unsupervised method designed to accurately estimate hidden factors (sources of variation) in single cell gene expression data while adjusting for known factors. The iterative and flexible framework of IA-SVA allows the accurate estimation of multiple and potentially correlated factors along with their statistical significance, which is the main advantage of IA-SVA over existing methods. This flexibility is more realistic given the confounded nature of known and unknown factors in single cell gene expression measurements. Therefore, IA-SVA has an improved performance over existing SVA-based methods in terms of estimating hidden sources of variation when they are correlated with each other and with known variables. IA-SVA is an effective alternative to methods developed for single cell data analyses (e.g., CellView and Seurat), especially for the analyses of complex data (i.e., data with multiple confounding and correlated factors). With the increasing amount of single cell studies and the increasing complexity of human cohorts, IA-SVA will serve as an effective statistical framework specifically designed to handle unique challenges of scRNA-seq data analyses.

## Methods

### IA-SVA framework

We model the log-transformed sequencing read counts for *m* cells and *n* genes (i.e.*Y*_*m* × *n*_) as a combination of known and unknown variables as follows:

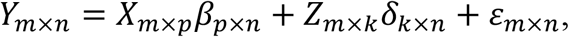

where *X*_*m*×*p*_ is a matrix for *p* known variable(s) (e.g., group assignment for cases and controls, sex or ethnicity), *Z*_*m*×*k*_ is a matrix for *k* unknown variables and ε_*m*×*n*_ is the error term. With this model, we can account for any clinical/experimental information about samples (e.g., sex, ethnicity, age, BMI or batch) as known factors (*X*_*m*×*p*_) and dissect unaccounted variation in the read count data that is attributable to hidden factors (*Z*_*m*×*k*_).

Existing unsupervised SVA-based methods (e.g., USVA^10^, RUV^13^, ISVA^12^) obtain the residual matrix (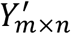) by regressing read counts (*Y*_*m*×*n*_) on all known factors (*X*_*m*×*p*_). Then, they infer the hidden factors from this residual matrix (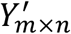) using dimensionality reduction algorithms (e.g., PCA, SVD or ICA). Thus, by definition, multiple hidden factors captured by these methods are orthogonal to each other and to known variables. Therefore, if hidden factors are correlated with each other and with known factors, the direct inference from the residual matrix leads to biased estimates of hidden factors due to the orthogonality assumption.

In contrast, IA-SVA does not impose orthogonality between factors (hidden or known) and allows an unbiased estimation of correlated factors via a novel iterative framework (Figure 1). At each iteration, IA-SVA first obtains residual matrix (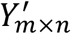), i.e., read counts adjusted for all known factors (*X*_*m*×*p*_) including surrogate variables of unknown factors estimated from previous iterations and extracts the first principal component (PC1) from the residuals (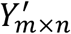) using SVD. Next it tests the significance of PC1 in terms of its contribution to the unmodeled variation (i.e., residual variance). Using this PC1 as a surrogate variable (as in the case of existing methods) implicitly imposes orthogonality between known and hidden factors. Instead, IA-SVA uses PC1 to infer gene weights, which are also used to infer genes associated with the hidden factor. IA-SVA relies on the fact that the first principal component (PC1) of the residual matrix is highly correlated with the hidden factor that contributes the most to the unmodeled variation in data, and thus, PC1 can be used to sort genes in terms of their relative association strength with the hidden factor. To infer these genes, IA-SVA regresses *Y* on PC1 and calculates the coefficient of determination (*R^2^*) for each gene. Genes with high *R^2^* scores can be treated as marker genes for the factor. These *R^2^* scores are further utilized for an unbiased inference of the hidden factor while retaining the correlation structure between known and hidden factors. For this, IA-SVA first obtains a weighted read count matrix (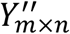) by weighing all genes with respect to their *R^2^* scores (i.e., 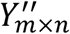 = *Y*_*m*×*n*_ *W*_*n*×*n*_ where *W* is a diagonal matrix of *R^2^* values). Then it conducts a SVD on 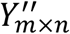 and obtains the PC1 to be used as a surrogate variable (SV) for the hidden factor. In the next iteration, IA-SVA uses this SV as an additional known factor to identify further significant hidden factors. The iterative procedure of IA-SVA is composed of six major steps as summarized in Figure 1A and below:

**[Step 1]** Regress *Y*_*m*×*n*_ on all known factors (*X*_*m*×*p*_), including SVs obtained from previous iterations, to obtain residuals (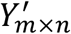).

**[Step 2]** Conduct a SVD on the obtained residuals (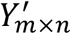) to extract the first PC (PC1).

**[Step 3]** Test the significance of the contribution of PC1 to the variation in residuals (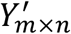) using a non-parametric permutation-based assessment ^9,10,29^ as explained further in the next section.

**[Step 4]** If PC1 is significant, regress *Y*_*m*×*n*_ (in this case do not use the residual matrix to be able to capture factors correlated with known factors) on PC1 to compute the coefficient of determination (*R^2^*) for every gene. If PC1 is not significant, stop the iteration and conduct subsequent down stream analysis using previously obtained significant SVs.

**[Step 5]** Weigh each gene in *Y*_*m×n*_ with respect to its *R^2^* value by multiplying a gene’s read counts (*Y*_*m*×*n*_) with its *R^2^* values (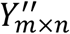 = *Y*_*m*×*n*_*W*_*n*×*n*_). The highly weighted genes in this framework serve as the genes affected by the hidden factor.

**[Step 6]** Conduct a second SVD on this weighted read counts matrix (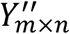 = *Y*_*m*×*n*_*W*_*n*×*n*_)

obtain PC1, which will be used as the surrogate variable (SV) for the hidden factor.

At the end of this six-step procedure, IA-SVA uses the detected SV (if significant) as an additional known factor in the next iteration. The algorithm stops, when no more significant PC1s are detected in Step 3. Significant SVs obtained via IA-SVA can be used in subsequent analyses. If an SV arises from an unwanted factor (e.g., cell contamination), these SVs can be included as covariates in the model to remove the unwanted variation or to filter out contaminated cells. In single cell data significant SVs could also explain ‘wanted’ biological factors (e.g., different cell types) and genes associated with such SVs can be further evaluated to discover novel biology from these complex datasets.

### Assessing the significance of hidden factors

To test the significance of the contribution of a hidden factor estimate (i.e., PC1 obtained in Step 2) to the residual variation, we used the permutation based significance test as previously applied in the surrogate variable analysis ^10,29^. Unlike SVA ^10^, which tests all putative hidden factors at once, IA-SVA assesses the significance of hidden factors one at a time during the corresponding iteration (always for the PC1 detected in that iteration). Briefly, IA-SVA i) conducts a SVD on the residual matrix obtained from Step 1, ii) computes the proportion of variation in this matrix explained by the first singular vector (i.e., PC1) and iii) compares this proportion against the values obtained from permuted residual matrices, as further explained below:

**[Step 1]** Conduct a SVD on the residual matrix (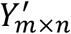).

**[Step 2]** Calculate the proportion of residual variance explained by the first singular vector (PC1) using the test statistic: 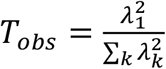, where *λ*_*k*_ is the *k*-th singular value.

**[Step 3]** Generate a permuted residual matrix by i) permuting each row of the log-transformed read count matrix *Y*_*m*×*n*_ and regressing the permuted read count matrix on all known factors (*X*_*m*×*p*_) to obtain fitted residuals.

**[Step 4]** Repeat Step 3 *M* times and generate an empirical null distribution of the test statistic by calculating (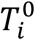, *i* = 1, …, *M*) for the *M* permuted residual matrices.

**[Step 5]** Compute the empirical p-value for the first singular vector (PC1) by counting the number of times the null statistics (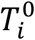) exceeds the observed one (*T_obs_*) divided by the number of permutations (*M*).

### scRNA-seq data simulation

To eliminate the potential bias in data simulations and make simulation studies more objective ^30^, we used a third-party simulation software (Polyester R package ^31^) and study design (http://jtleek.com/svaseq/simulateData.html) and simulated scRNA-seq data to test IA-SVA’s performance. The original simulation design is slightly modified to reflect characteristics of scRNA-seq data for high dropout rate (i.e., excessive number of zeros in the data) and multiple hidden factors highly correlated with known factors. First, to simulate high dropout rates (proportion of zero counts = ~70%), we estimated Polyster’s zero-inflated negative binomial model parameters (i.e., p_0_: probabilities that the count will be zero, mu: mean of the negative binomial, size: size of the negative binomial) from real-world scRNA-seq data from human pancreatic islets using the Fludigm’s C1 platform ^17^. Using these estimated model parameters, we simulated expression data for *m* cells and *n* genes under two hypotheses: 1) the null hypothesis: no hidden sources of variation, and 2) the alternative hypothesis: three hidden factors with two values (−1 vs. 1). Under both scenarios, we simulated a primary variable of interest (i.e., case vs. control) and simulated 10% of genes to be differentially expressed between the two groups. Under the alternative hypothesis, we simulated three hidden factors that affect 30%, 20% and 10% of randomly chosen genes respectively and simulated two different scenarios where these factors are moderately correlated (|*r*|*=~*0.3-0.6) or weakly correlated (|*r*|*<*0.3) with the group variable.

### Data processing and normalization

In all analyses, we filtered out low-expressed genes with read counts <= 5 in less than three cells and normalized the retained gene expression counts using SCnorm^19^ with default settings for further analyses. For single cell data visualization examples, we normalized generead counts by dividing each cell column by its total counts then multiplying median of library size, which is similar to the default normalization method “LogNormalize” implemented in Seurat ^16^.

### Availability of data and methods

An R package for IA-SVA with example case scenarios is freely available from https://github.com/UcarLab/IA-SVA. The published data sets analyzed in this study including single-cell RNA sequencing read counts and annotations describing samples and experiment settings are included in an R data package (iasvaExamples) deposited at https://github.com/dleelab/iasvaExamples.

## Acknowledgements

We thank the Jackson Laboratory Computational Science group, Ucar and Stitzel lab members for constructive feedback throughout this project. We thank Jane Cha, JAX scientific illustrator, for her help with Figure 1. This study was made possible by generous financial support of the National Institute of General Medical Sciences (NIGMS) under award number GM124922 (to DU) and by the Jackson Laboratory Scientific Services Innovation Fund (to D.L. and D.U.) Opinions, interpretations, conclusions, and recommendations are solely the responsibility of the authors and do not necessarily represent the official views of the National Institutes of Health (NIH).

## Author Contributions

D.L. and D.U. designed the project, generated the figures and wrote the manuscript. D.L. developed the statistical framework and run the data analyses. M.B. provided advice on data analyses and interpretation of results. A.C. contributed to the data pre-processing and the generation of the R package. All authors read and approved this manuscript prior to submission.

## Additional Information

Supplementary information accompanies this paper at http://www.nature.com/srep. Competing financial interests: The authors declare no competing financial interests.

